# Common viral infections seed regionally distinct resident memory T cells in the human CNS

**DOI:** 10.64898/2026.09.21.753190

**Authors:** Hanna N. Degefu, Tyler G. Searles, Tiffany Chen, Shawn C. Musial, Jordan F. Isaacs, Sierra A. Kleist, Shizhao Yang, Myles Ford, Fred W. Kolling, Mary Jo Turk, Alexander G J Skorput, Chun-Chieh Lin, George J. Zanazzi, Jennifer Hong, Li Song, Pamela C. Rosato

**Affiliations:** Department of Microbiology and Immunology, Geisel School of Medicine at Dartmouth College, Lebanon, NH; Department of Neurology, Dartmouth Health, Lebanon, NH; Dartmouth Cancer Center, Lebanon, NH; Department of Pathology and Laboratory Medicine, Dartmouth Health, Lebanon, NH; Department of Surgery, Dartmouth Health, Lebanon, NH; Department of Biomedical Data Science, Dartmouth College, Lebanon NH

## Abstract

T cells persist in the central nervous system (CNS) and can drive both protection and neurological disease. How these cells are organized in humans and what they recognize is largely unknown. Here, we profiled CD8□ T cells across anatomically distinct CNS regions, obtained through on-site autopsies and temporal lobe resection surgeries, using single-cell RNA sequencing, paired T cell receptor sequencing, and DNA-barcoded tetramers. Resident memory T cells (T_RM_) specific for Epstein-Barr virus, cytomegalovirus, influenza A, and SARS-CoV-2 were identified across CNS compartments. Anatomical location was the strongest correlate of T_RM_ cell state, with leptomeningeal cells adopting a cytokine-poised T_RM_ program, whereas brain T_RM_ cells were transcriptionally restrained. Cells of the same clonotype spanned tissues yet adopted local transcriptional states. Viral specificity added another layer of T_RM_ heterogeneity with *GZMK/GZMA-*expressing EBV-specific populations and interferon-stimulated gene signatures in SARS-CoV-2 and Influenza A-specific cells. The human CNS thus harbors regionally distinct CD8^+^ T_RM_ shaped by common viral exposures.

## INTRODUCTION

The central nervous system (CNS) has historically been viewed as an immune-privileged site, but it is now recognized as an immunologically surveilled tissue in which resident and recruited immune cells shape host defense, tissue homeostasis and disease^1–4^. T cells are present across multiple human CNS compartments, including the brain parenchyma^5–8^, meninges^7–9^ and cerebrospinal fluid^8,10,11^, where they display features of tissue-resident memory T cells (T_RM_), including expression of tissue-retention molecules and reduced expression of recirculating molecules. CD8^+^ T_RM_ are long-lived, non-recirculating sentinels that provide rapid local protection after antigen re-encounter in tissues throughout the body^12–14^. Despite this well-defined role in peripheral tissues, the organization, antigen specificity and functional state of these cells in the CNS, particularly in humans, remains incompletely understood.

Much of what is known about CNS T_RM_ derives from mouse models. Brain T_RM_ arise not only after direct neurotropic infection^15–20^ but also following peripheral infections^21–24^, vaccinations^21,25^, gut microbiota colonization^26^, physiologic microbial exposure^27^, and even intravenous T cell transfer models devoid of antigen and neuroinflammation^23,28,29^. Once established, they mediate rapid local protection against reinfection through cytokine production and cytotoxicity^15,18,21,30^, while in other contexts they contribute to blood brain barrier disruption, neuroinflammation and tissue injury^19,20,31,32^. Notably, peripherally induced brain T_RM_ (i.e. via vaccination or peripheral infections) in mice accumulate with successive exposures and distribute across anatomically distinct CNS regions^22^, suggesting that the CNS T_RM_ compartment reflects cumulative systemic immune history and may contain clonally expanded populations specific for common peripheral pathogens. Whether these principles extend to the human CNS, however, is unclear.

Studying CNS T_RM_ in humans has been difficult; while meaningfully present, the abundance of CNS T cells is roughly about 200-fold below barrier tissues^29,33^. Moreover fresh, viable specimens spanning multiple compartments of the same individual are difficult to obtain. CNS regions also differ in both function and local environment. For example, in the brain, the frontal and temporal lobes support cognition, language and memory, the hippocampus is central to learning and memory, and the cerebellum coordinates movement, while the surrounding meninges form a protective border, with lymphatics in the dura^34–38^. These regions contain distinct mixtures of specialized cells such as neurons and glia in the brain, and fibroblasts and pericytes in the meninges, creating local environments that may shape immune cell adaptation^39^. Indeed, a recent study profiling T cells from postmortem CNS tissues from donors with and without neurologic disease showed by flow cytometry that distinct anatomic sites harbor T cells with different phenotypes^8^. It is unclear if this data reflects distinct T cell specificities or clones in different anatomic compartments, or if clonally identical T cells are differentially shaped by their tissue environment. Another recent report showed that the leptomeninges harbor clonally expanded CD8^+^ T_RM_ whose repertoires overlap with the brain^7^. These important studies give us insights into the T cell landscape of the human CNS, but how T_RM_ are organized across CNS regions, and whether their identity and clonal relationships are shaped by anatomical location or antigen specificity is still unclear.

The antigens recognized by human CNS T cells are perhaps least understood. In neurodegenerative diseases and epilepsy, expanded CNS-associated clones have been linked to common viruses, including cytomegalovirus (CMV), influenza A virus (IAV) and Epstein–Barr virus (EBV)^7,40,41^. However, antigen specificity was often inferred computationally from public TCR– antigen databases that are incomplete and biased toward well-characterized viral epitopes. Direct, peptide-MHC multimer identification of EBV-specific T cells in the human CNS has also been reported in tumor tissue and multiple sclerosis (MS)^42–44^, however, whether virus-specific CNS T_RM_ are broadly present across individuals remains unknown. Persistent and recurrent viral infections are strong candidates to seed the CNS T_RM_ pool. For example, herpesviruses such as EBV and CMV establish lifelong latency and drive large, durable CD8^+^ responses^45,46^; IAV causes repeated infections that generate tissue-localized memory^47^; and SARS-CoV-2 generates host-wide T cell immunity and is linked to neurological sequela^48^. Whether such systemic exposures seed antigen-specific CD8□ T_RM_ across the human CNS, and whether different specificities adopt distinct clonal and transcriptional programs, is unknown.

Here, we combine multiparameter flow cytometry, single-cell RNA sequencing, paired TCR sequencing and DNA-barcoded peptide-MHC tetramer profiling across postmortem multi-region CNS samples and fresh surgical resections with matched blood to define the organization and antigen specificity of human CNS CD8^+^ T cells. We show that CNS CD8^+^ T cells adopt distinct tissue-resident programs correlated with anatomic compartment, are oligoclonally expanded and include dominant clonotypes shared across tissue regions. Using HLA-A*02 tetramers, we directly identify CD8^+^ T_RM_ specific for EBV, CMV, IAV and COVID in cerebellum, hippocampus, frontal lobe, temporal lobe and leptomeninges, revealing antigen-dependent patterns of clonal expansion, tissue sharing and distinct phenotypes that reflected tissue of origin and viral specificity. Together, these findings demonstrate that common viral exposures shape a clonally connected, antigenically diverse CD8^+^ T_RM_ network in the human CNS, reframing CNS immunosurveillance as an active process that reflects an individual’s infection history.

## RESULTS

### Virus-specific CD8^+^ T cells are present across human CNS compartments

The T cell landscape of the human CNS has been difficult to define, in large part because capturing antigen-specific T cells at a meaningful resolution requires substantial amounts of fresh, viable tissue that are rarely accessible. While recent studies have begun to characterize CD8^+^ T cells in select CNS compartments or specific disease contexts^7,8,26,41,43^, a matched multi-region profiling that resolves T cell phenotype, clonal relationships and antigen-specificity within the same donor has not been achieved. As such, a central unresolved question is whether CD8^+^ T cell identity, migratory status, and antigen specificity vary across anatomically distinct CNS regions, and which antigens these cells recognize. To address this, we established an on-site autopsy pipeline that enables dissection and processing of different CNS compartments, including leptomeninges and frontal lobe, from individual donors, together with matched lung tissue as a peripheral non-lymphoid benchmark for direct comparison with CNS CD8□T cells.

We reproducibly isolated viable CD8□T cells from all sampled regions within individual autopsies for flow cytometric analysis (**Fig. 1a**). Most CD8^+^ T cells exhibited a CCR7□CD45RA□ phenotype, with smaller CCR7□CD45RA^+^ populations and few CCR7^+^CD45RA^+^ (naïve) or CCR7^+^CD45RA^-^cells, consistent with previous studies^6,8,49^ (**Supp Data Fig. 1a)**. Across tissues, CD69□cells predominated, and CNS CD8□T cells displayed CD69^+^CD103^+^ profiles consistent with tissue residency, with frontal lobe and leptomeninges showing comparable distributions (**Fig. 1b**), consistent with prior reports of CD8^+^ T_RM_ in human CNS^6–8^.

**Figure 1.**
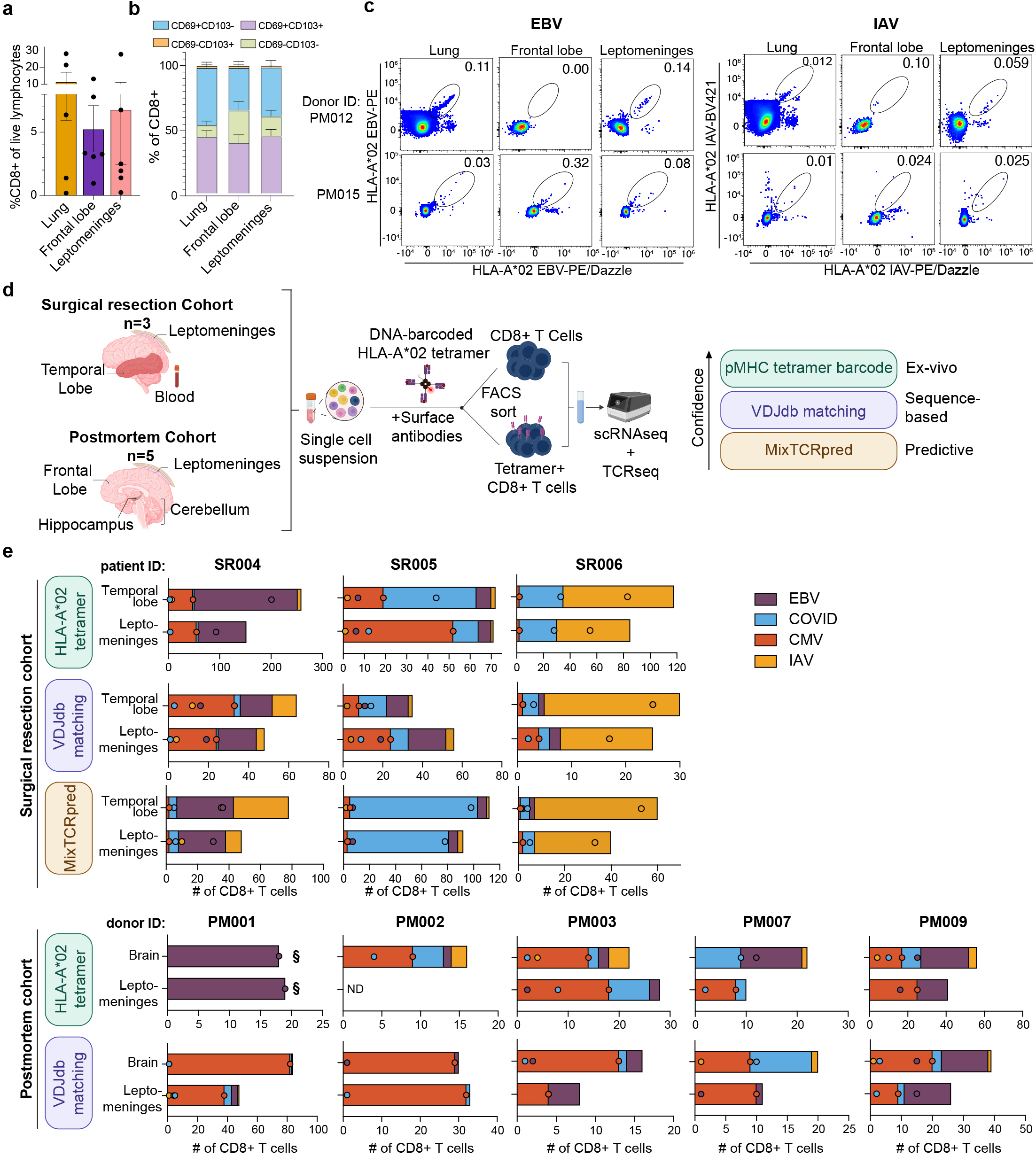
Virus-specific CD8^+^ T cells are present across human CNS compartments. (**a**) Frequency of CD8^+^ T cells among live lymphocytes in human lung, brain frontal lobe and leptomeninges. (**b**) CD69 and CD103 co-expression among CD8^+^ T cells across tissues (n=5; bars represent mean ± SEM). (**c**) Flow cytometry of combinatorial dual tetramer staining with Epstein Barr virus (EBV, left) and influenza A virus (IAV, right) HLA-A*02 tetramers from two donors (rows). Gated on live CD3^+^ CD8^+^ CD4^-^T cells negative for background single fluorophore staining and irrelevant tetramer fluorophores. (**d**) Schematic of post-mortem (n=5) and surgical resection (n=3) cohorts, experimental workflow, and hierarchy of antigen-specific T cell identification methods stratified by confidence level. (**e**) Number of virus-specific CD8+ T cells recovered from the leptomeninges and brain tissue for each donor, stratified by identification method and donor cohort where SR=surgical resection (top) and PM=postmortem (bottom). Brain tissue for the PM cohort encompasses frontal lobe, hippocampus, and cerebellum. Dots represent the absolute number of T cells per specificity, and stacked bars are the cumulative total per donor. All tissues were stained with HLA-A*02 tetramers to EBV, cytomegalovirus (CMV), IAV, and SARS-CoV-2 (COVID) with the exception of PM001 (§) which was stained for EBV tetramer only. ND= none detected. Full tissue-level counts provided in Supplementary Table 2.

We next asked whether CNS CD8^+^ T cells include cells specific for common viral infections. Recent studies in mice have shown that CNS T_RM_ can be established not only by neurotropic infections, but also by peripheral infection and vaccinations^21,22,27,50^. Given this, we hypothesized that a population of CNS CD8^+^ T cells is shaped by prior systemic viral exposures. To test this, we used EBV and IAV-specific HLA-A*02:01 peptide-MHC tetramers to stain CD8^+^ T cells from tissues from two donors with no significant neurological disease at the time of death **(Table 1**, PM012 and PM015). We detected tetramer-positive CD8^+^ T cells for EBV and IAV in the leptomeninges, and for EBV in the frontal lobe in one donor, providing proof-of-concept data that virus-specific CD8^+^ T cells can be present within CNS tissue and may not be restricted to peripheral sites (**Fig. 1c, Supp Data Fig. 1b**). These cells predominantly expressed CD69, with variable CD103 co-expression, consistent with a tissue resident phenotype (**Supp Data Fig. 1c**).

To extend these observations across more donors, and to include multiple viral specificities and anatomical regions at single-cell resolution, we established two independent cohorts (**Fig. 1d**). The surgical resection (SR, n=3) cohort provided fresh viable tissue from temporal lobe, leptomeninges, and matched peripheral blood from donors undergoing temporal lobe resection for drug-refractory epilepsy. The postmortem (PM, n=5) cohort comprised donors from in-house autopsies performed at Dartmouth Hitchcock Medical Center (as described above), which we expanded to sample cerebellum, frontal lobe, hippocampus, and leptomeninges. Together, these two cohorts provided complementary anatomical breadth, tissue volume, and tissue quality necessary to resolve questions that have been difficult in previous human CNS studies **(Table 1).** For both cohorts, isolated CD8^+^ T cells were stained with DNA-barcoded HLA-A*02:01 tetramers for immunodominant epitopes from EBV, CMV, IAV, and SARS-CoV-2 (COVID). For technical reasons, donor #PM001 was stained with EBV tetramer only. Tetramer+ CD8^+^ T cells were enriched by dual label flow sorting and then combined with the non-enriched CD8^+^ T cell fraction prior to paired single-cell RNA and TCR sequencing **(Fig. 1d)**. This approach allowed for simultaneous single cell resolution of antigen specificity, transcriptional phenotype, and clonal identity.

Tetramer staining identified virus-specific CD8^+^ T cells in the brain and leptomeninges across all eight donors from both cohorts (**Fig. 1e**). To complement this, we applied two TCR sequence-based approaches; MixTCRpred which is a machine learning-based epitope predictor using paired TCRα+TCRβ sequences^51^, and TCR sequence matching against the curated VDJ database (VDJdb) of known antigen-specific TCRs. As with direct tetramer staining, these approaches identified virus-specific CD8^+^ T cells in all donors across both cohorts (**Fig. 1e**). In the SR cohort, all three methods identified cells targeting each of the four viral specificities, and cross-method concordance analysis showed that MixTCRpred and VDJdb assignments corresponded to the viral epitopes defined by tetramer binding (**Supp Data Fig 1d; Table 2**). Thus, complementary experimental and sequence-based methods identified virus-specific CD8^+^ T cells in CNS tissues of all donors across both cohorts.

Of note, fewer tetramer-positive cells and paired TCRα–TCRβ sequences were recovered from PM than from SR samples, likely reflecting differences in tissue handling: PM tissues were processed up to 32 h after death, whereas SR tissues were processed fresh immediately after resection in the operating room. Lower viability and postmortem degradation may have reduced TCR recovery and impaired tetramer binding^52^. Consequently, MixTCRpred assignments were limited in PM samples, whereas VDJdb matching was less affected because it permitted TCRβ-only matches, which constituted most matches in the PM cohort (**Table 2**).

We next examined how viral specificities varied among donors. Virus-specific CD8^+^ T cells targeting all four specificities were recovered from both cohorts but the relative contribution of each specificity varied across donors, with CMV enriched in older postmortem donors (PM001, 002, 003 and 009, ≥90yo) while EBV and CMV frequencies were lower in the youngest surgical resection donor (SR006, 20yo). VDJdb matching additionally returned low-frequency matches to other viral specificities including dengue virus, HCV, and HIV-1 **(Table 3).** Together, these data provide direct evidence that common peripheral viral infections broadly seed antigen-specific CD8^+^ T cells across human CNS compartments.

### Virus-specific CD8 T cells adopt tissue-resident transcriptional programs in the CNS

Based on our flow phenotyping data **(Fig 1)**, and previously published reports of T_RM_ in the CNS^5–11^, we hypothesized that the identified virus-specific CD8^+^ T cells were largely resident rather than blood contaminants or recirculating populations. To assess this, we performed dimensionality reduction and unsupervised clustering on the single-cell RNA sequencing data from both cohorts. In the PM cohort, tissue of origin emerged as a major driver of transcriptional variation, with leptomeningeal cells segregating into distinct regions of the UMAP while brain-derived cells from cerebellum, hippocampus and frontal lobe showed greater overlap (**Fig. 2a**). A similar pattern was observed for the SR cohort, where leptomeninges and temporal lobe formed largely non-overlapping transcriptional spaces separate from blood-derived cells. These patterns were consistent across individual donors (**Supp Data Fig. 2a**). Unsupervised clustering identified 13 transcriptionally distinct CD8^+^ T cell populations in each cohort (**Fig. 2b**). To distinguish tissue-resident from circulating transcriptional states, we scored each cell using a T_RM_ gene signature adapted from Burn et. al.^53^ and calculated a UCell residency index per cell to classify clusters as T_RM_ or circulating (T_CIRC_) in each cohort (**Fig. 2c**). Across both cohorts, the majority of CNS CD8^+^ T cell clusters were classified as T_RM_ based on their residency index scores. As expected, expression of canonical residency-associated genes including *CD69, ITGAE* and *ZNF683* were enriched in T_RM_ clusters while circulating clusters showed higher expression of *S1PR1* and *SELL*, supporting subset classification (**Fig. 2d**).

**Figure 2.**
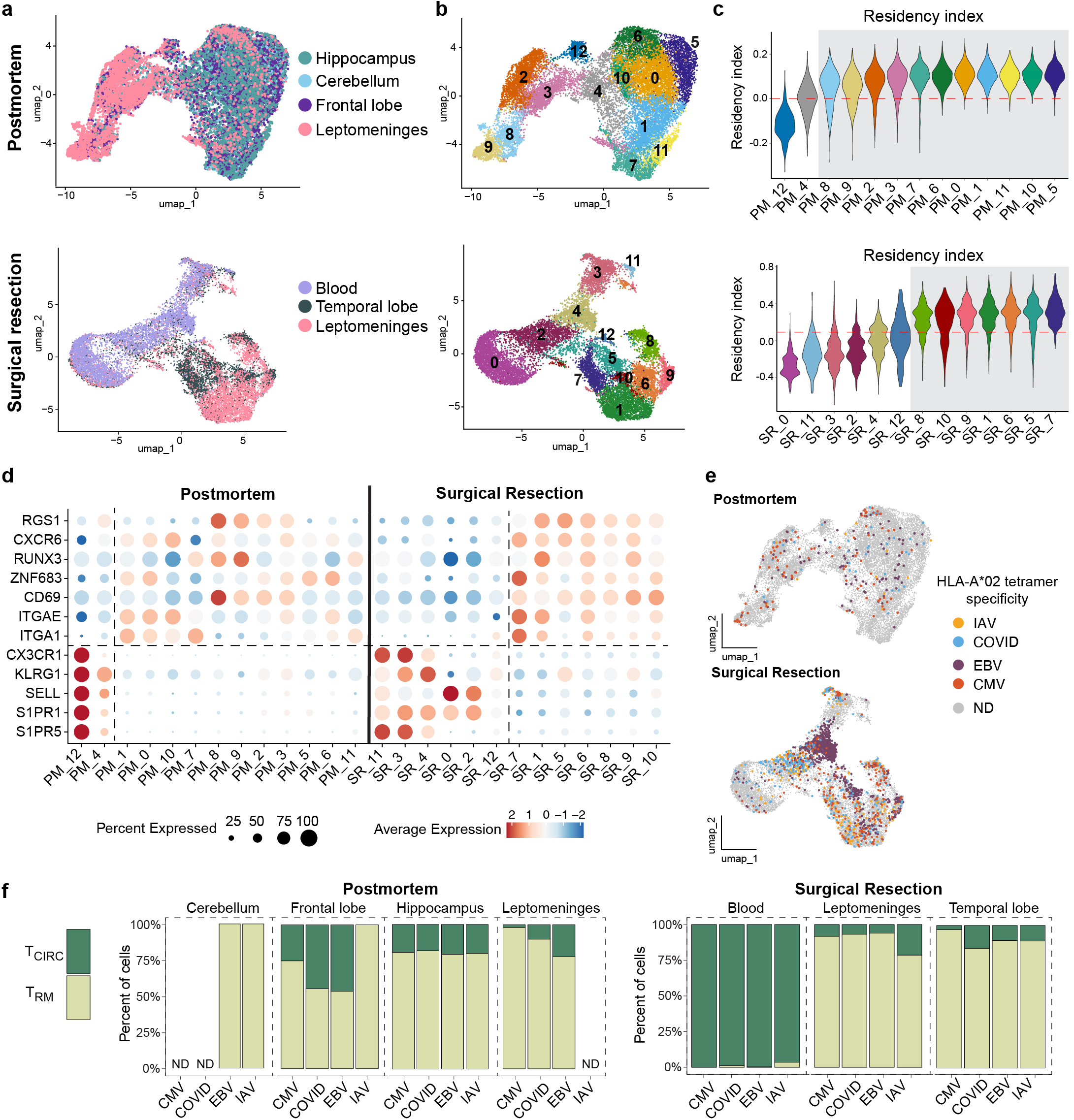
Virus-specific CD8^+^ T cells adopt tissue-resident transcriptional programs in the CNS. UMAP of CD8^+^ T cells from the postmortem (PM, n=5, top) and surgical resection (SR, n=3, bottom) cohorts annotated by tissue of origin (**a**) and unsupervised cluster identity (**b**). **(c**) UCell residency index distribution by cluster, calculated as the difference between T_RM_-upregulated and T_RM_-downregulated gene signature scores adapted from Burn et al.^44^ Dashed line indicates zero (PM, top), or a circulating threshold defined using blood-derived clusters as a transcriptional reference (SR, bottom). Clusters at or above the dashed line were classified as resident (T_RM_, gray box) and those at or below zero as circulating (T_CIRC_). (**d**) Dot plot showing scaled expression of select residency-and circulating-associated genes across clusters from both cohorts. Dot size represents percent of cells expressing each gene; color represents mean scaled expression. (**e**) UMAP of PM (top) and SR (bottom) CD8^+^ T cells with tetramer-identified virus-specific cells annotated by viral specificity. (**f**) Proportion of virus-specific CD8^+^ T cells assigned to T_RM_ or T_CIRC_ clusters, stratified by tissue and viral specificity. ND= not detected.

Having established the migratory landscape of total CNS CD8^+^ T cells, we next asked whether tetramer+ virus-specific cells preferentially adopt T_RM_ transcriptional states. Across CNS compartments in both cohorts, virus-specific CD8^+^ T cells of all four specificities were largely found within T_RM_ clusters, with blood-derived virus-specific cells showing the expected enrichment in T_CIRC_ clusters (**Fig. 2e,f**). These findings demonstrate that the majority of virus-specific CNS CD8^+^ T cells are not simply blood contaminants or recirculating cells, but instead largely adopt tissue-resident transcriptional programs within brain and leptomeningeal compartments, regardless of antigen specificity.

### Anatomic location is the primary determinant of CNS CD8^+^ T_RM_ identity

Given the segregation of clusters by tissue of origin (**Fig 2**), we next tested if CD8^+^ T_RM_ adopt distinct cell states across the different CNS regions. Focusing on defined T_RM_ clusters (T_CIRC_ clusters in **Supp Data Fig. 3a**), dot plot analysis of gene categories spanning stem/progenitor, killer cell lectin-like receptors (KLRs), cytotoxic, cytokine, exhaustion/inhibitory, and interferon-stimulated gene (ISG) programs revealed functional heterogeneity across clusters in both cohorts (**Fig. 3a-d**). Leptomeninges-enriched clusters consistently showed high expression of effector cytokine and chemokine transcripts including *IFNG, CCL4, CCL3,* and *TNF*. Because T cells isolated from these tissues do not produce cytokines at the protein level unless stimulated^6^ (**Supp Data Fig. 3b**), these transcripts likely represent a “poised” rather than an actively secreting state, consistent with T_RM_ described in the lung^54^. In contrast to leptomeninges, brain tissue clusters were transcriptionally diverse, encompassing populations defined by KLR gene expression including *KLRB1, KLRC1,* and *KLRD1*, stem/progenitor-associated genes including *SATB1* and *FOXN3*, high ISGs, and exhaustion/inhibitory markers including *TOX, LAG3,* and *HAVCR2*. Granzyme expression further delineated two distinct cytotoxic clusters: one defined by selective *GZMK* and *GZMA* expression with minimal *GZMB* or *GNLY* (PM cluster 10 and SR cluster 5), and another characterized by high *GZMB* and *GNLY* expression (PM C5). These transcriptional identities were largely consistent across independent cohorts, with pseudobulk correlation confirming conservation of major CD8^+^ T cell transcriptional states (**Supp Data Fig. 3c**).

**Figure 3.**
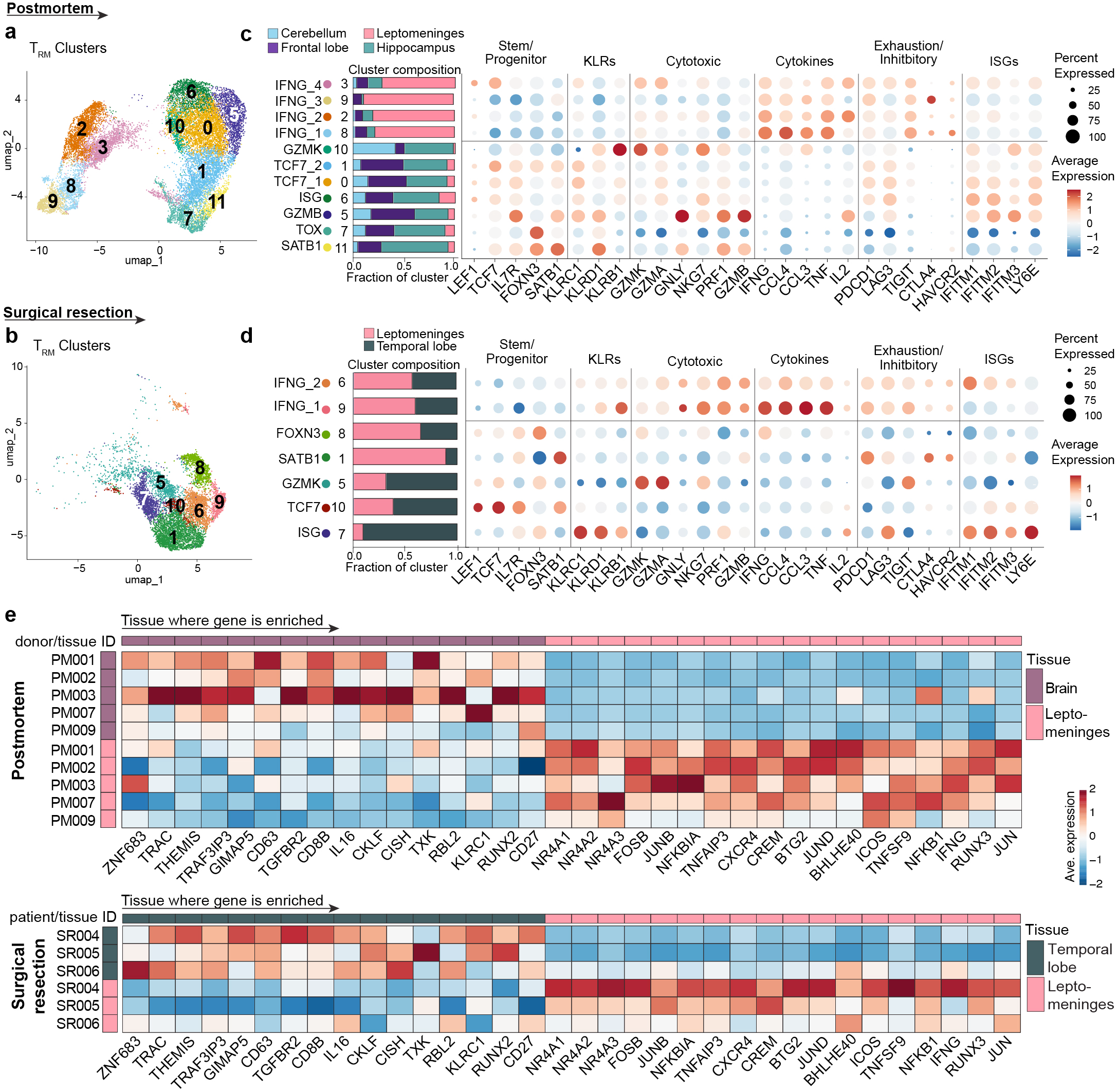
Anatomic location is the primary determinant of CNS CD8^+^ TRM identity. (**a and b**) UMAP of CD8^+^ TRM postmortem (PM, n=5, a) and surgical resection (SR, n=3, **b**) cohorts with T_CIRC_ clusters removed. (**c and d**) Dot plot showing scaled expression of selected gene categories across T_RM_ clusters in PM (**c**) and SR (**d**) cohorts. Rows annotated by cluster identity and composition indicated by stacked bar graphs representing the proportion of cells from each tissue. Dot size represents percent of cells expressing each gene; color represents mean scaled expression. ISGs=Interferon-stimulated genes, KLRs= Killer cell lectin-like receptors. (**e**) Heatmap showing z-score normalized average gene expression of a representative set of DEGs distinguishing leptomeningeal from brain tissue TRM in PM (top) and SR (bottom) cohorts. Color scale represents z-scored expression clipped to ±2.

Next, to identify a transcriptional signature distinguishing leptomeningeal from brain tissue T_RM_, we performed differential gene expression analysis comparing leptomeningeal and brain T cells independently for each cohort and identified overlapping differentially expressed genes (DEGs) reproducible across both datasets. Visualizing a representative set of reproducible DEGs across both cohorts, brain-enriched genes reflected a canonical T_RM_ program including *ZNF683* (Hobit), quiescence regulators such as *GIMAP5*, *CISH*, and TGF-β receptor type 2 (*TGFBR2*), while leptomeninges-enriched genes included NR4A family transcription factors, AP-1 components, NF-κB pathway genes, and effector cytokine transcripts *IFNG* and *CCL4* (**Fig. 3e**). Because leptomeninges and brain tissue underwent collagenase digestion for different durations, and because AP-1 family genes (e.g. *FOS*, *JUN*) have been previously associated with digestion artifacts^55^, we repeated psuedobulk PCA after the removal of published T cell specific digestion-associated genes^56^ (**Supp Data Fig. 3d, e**). Samples continued to separate by tissue, indicating that transcriptional differences between leptomeningeal and brain T cells were not driven by differences in dissociation protocols. Overall, these findings show that tissue location is a primary determinant of CNS CD8^+^ T_RM_ identity.

### Clonally expanded CD8^+^ T_RM_ are shared across CNS tissues but adopt region-specific transcriptional states

We next used TCR clonal analysis to ask whether the tissue-associated T_RM_ states comprise distinct local clones or clonally related cells distributed across regions. We analyzed paired TCRα and TCRβ sequences data from both cohorts across all clusters. Clonotypes were defined within each donor by identical V and J gene usage and CDR3 nucleotide sequences. Mapping clonal expansion onto the transcriptional UMAP revealed that, in both cohorts, expanded clonotypes were enriched within T_RM_ clusters while circulating clusters were largely composed of singletons and small clones (**Fig. 4a**). Clone size distributions confirmed expanded clonotypes across all CNS compartments indicating that clonal expansion is a prominent feature of CNS CD8□T cells, consistent with a recent study^7^ (**Fig. 4b**). Donor level profiles further showed substantial heterogeneity in the magnitude of clonal expansion (**Supp Data Fig. 4a**).

**Figure 4.**
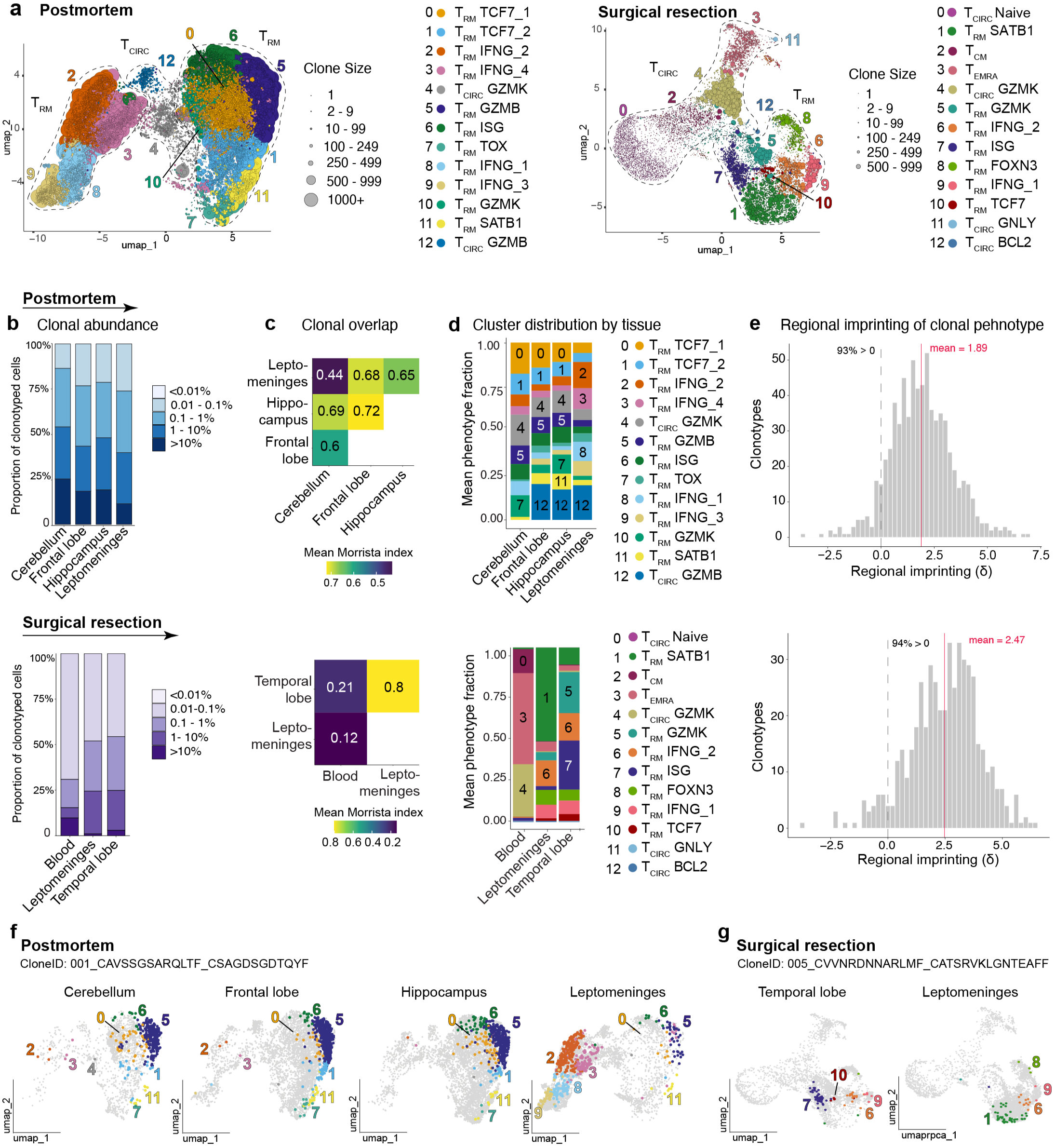
Clonally expanded CD8^+^ T_RM_ are shared across CNS tissues but adopt region specific transcriptional states. (**a**) UMAP of CD8+ T cells from the postmortem (PM, n=5, left) and surgical resection (SR, n=3, right) cohorts with dot size scaled to T cell clonotype size. Annotated by cluster identity. (b) Mean proportion of clonotyped CD8+ T cells across donor normalized clonotype abundance bins, averaged across donors with each tissue available; PM (top), SR (bottom). (**c**) Morisita overlap index heatmap showing pairwise clonal similarity between tissues in the PM cohort (top) and SR cohort (bottom). Values represent the mean Morisita index across donors. (**d**) Phenotypic composition of the top 10 expanded clonotypes per tissue averaged across donors. Colors represent cluster identity, top clusters per tissues annotated with cluster number. PM (top), SR (bottom). (**e**) Regional imprinting of shared clonotypes in brain and leptomeninges. Per cell, polarization score equals the z-scored brain UCell score minus the z-scored leptomeninges UCell score. Per clonotype, imprinting equals the mean (brain) minus the mean (leptomeninges). Tissue signatures were derived from cells belonging to tissue-restricted clonotypes. Dashed line at 0 marks no tissue imprinting of clonal phenotypes; red line = mean. PM cohort (top, n=565 clones) and SR cohort, excluding blood (bottom, n=455 clones). (**f and g**) UMAP of a representative T cell clone per donor: PM (donor ID: PM001) (**f**) and SR (donor ID: SR005) (**g**) faceted by tissue, illustrating transcriptional variability across CNS compartments. Cells belonging to the indicated clonotype are highlighted; all other cells shown in gray.

To test whether expanded clonotypes were restricted to individual CNS regions or shared across anatomic compartments we performed a Morisita overlap analysis, which revealed broad clonal sharing across CNS compartments in both cohorts (**Fig. 4c**). All CNS tissue pairs in the PM cohort showed moderate to high overlap (0.60-0.72), with the exception of cerebellum which had the lowest overlap with leptomeninges (0.43). In the SR cohort, temporal lobe and leptomeninges showed high overlap (0.80), while blood-brain overlap was substantially lower (0.14-0.22), indicating that expanded CNS clonotypes are largely distinct from the circulating repertoire. Jaccard indices and clonotype-level overlap confirmed consistent sharing patterns across tissue pairs in both cohorts (**Supp Data Fig. 4b,c**). Alluvial plots of the top 10 expanded clonotypes per donor also confirmed that the most expanded clones were largely shared across tissue regions within individual donors (**Supp Data Fig. 4d**). Despite broad clonal sharing, the phenotypic composition of top expanded clones was regionally distinct (**Fig. 4d**). In the PM cohort, leptomeningeal top clones were dominated by *IFNG*-expressing T_RM_ clusters (C2,3), while brain (cerebellum, frontal lobe and hippocampus) top clones were enriched for *TCF7, GZMB, GZMK*-expressing populations (C0, 1, 4, 5). In the SR cohort, leptomeningeal top clones were dominated by *SATB1*-expressing T_RM_ (C1) with secondary *IFNG* contributions (C6), while temporal lobe top clones were enriched for *GZMK*, ISG, and *IFNG*-expressing states (C5, 6, 7).

Finally, we asked whether shared clonotypes maintained a fixed transcriptional identity across tissues or instead, adopted region-specific states. For each clonotype present in both the brain and leptomeninges, we quantified a regional imprinting score, defined as the difference in polarization between the two tissue compartments within that clonotype, so that each clone served as its own control. Imprinting scores were positive for the large majority of shared clonotypes in both cohorts and were consistent across all donors, indicating that even T cell clones shared between tissues adopted the defined tissue-associated phenotypes (**Fig. 4e**, **Table 4**). Within shared clonotypes, both compartments largely matched their expected states, with brain T cells brain-leaning and leptomeningeal T cells leptomeninges-leaning (**Supp Data Fig. 4e**). Importantly, because the tissue polarization signature was derived from clonotypes restricted to a single tissue region so that the tested tissue-spanning shared clonotypes did not contribute to their own signature, and the effect was independent of sampling depth (**Supp Fig. 4f**). Representative examples show this pattern directly; in the PM cohort, T cells from a single expanded clone fell predominantly in the T_RM_-GZMB cluster (C5) across cerebellum, frontal lobe, and hippocampus. However, T cells from the same clone fell predominantly in T_RM_-IFNG clusters (C2, 3, 8 and 9) in the leptomeninges (**Fig. 4f**). Likewise in the SR cohort, a clone enriched in T_RM_-ISG (C7) in temporal lobe occupied T_RM_-SATB1 (C1) and T_RM_-IFNG clusters (C6 and 9) in the leptomeninges (**Fig. 4g**). Together, these data demonstrate that the human CNS contains a clonally expanded CD8□ T cell repertoire in which cells from the same clonotype are distributed across regions yet are transcriptionally imprinted by local tissue.

### Viral specificity shapes the clonal architecture and phenotype of antigen-specific CNS T_RM_

Having established that CNS CD8□ T cells are clonally expanded and regionally shared, we next asked whether antigen specificity influences the clonal organization and transcriptional phenotype of CNS T_RM_ or whether tissue region remains the dominant organizer regardless of specificity. Focusing on the SR cohort, which contained sufficient tetramer+ cells for cross-tissue clonal analysis, clonal size distribution analysis showed specificity-dependent differences in the magnitude of clonal expansion (**Fig. 5a**). EBV-specific T cells showed the highest proportion of expanded clonotypes, with the majority of cells belonging to clones of 500 cells or more whereas COVID-, CMV-, and IAV-specific cells were largely composed of singletons and small clones **(Fig. 5a)**. Clonal-sharing across compartments followed a similar trend (**Fig. 5b,c)**. While sharing across all three tissue compartments was limited, EBV-specific clonotypes showed the most extensive sharing, with 7.1% present across blood, temporal lobe and leptomeninges, consistent with some clonal distribution across both the CNS and circulation. In contrast, CMV-specific cells were largely compartment-restricted, with the largest fractions confined to the leptomeninges (52%) or temporal lobe (32%), and one clone dominant in blood but with limited CNS representation. COVID-and IAV-specific clonotypes were also predominantly blood restricted (45% and 43%) but lacked a dominant clone, and comprised multiple modestly expanded clones distributed across compartments **(Fig 5b,c)**.

**Figure 5.**
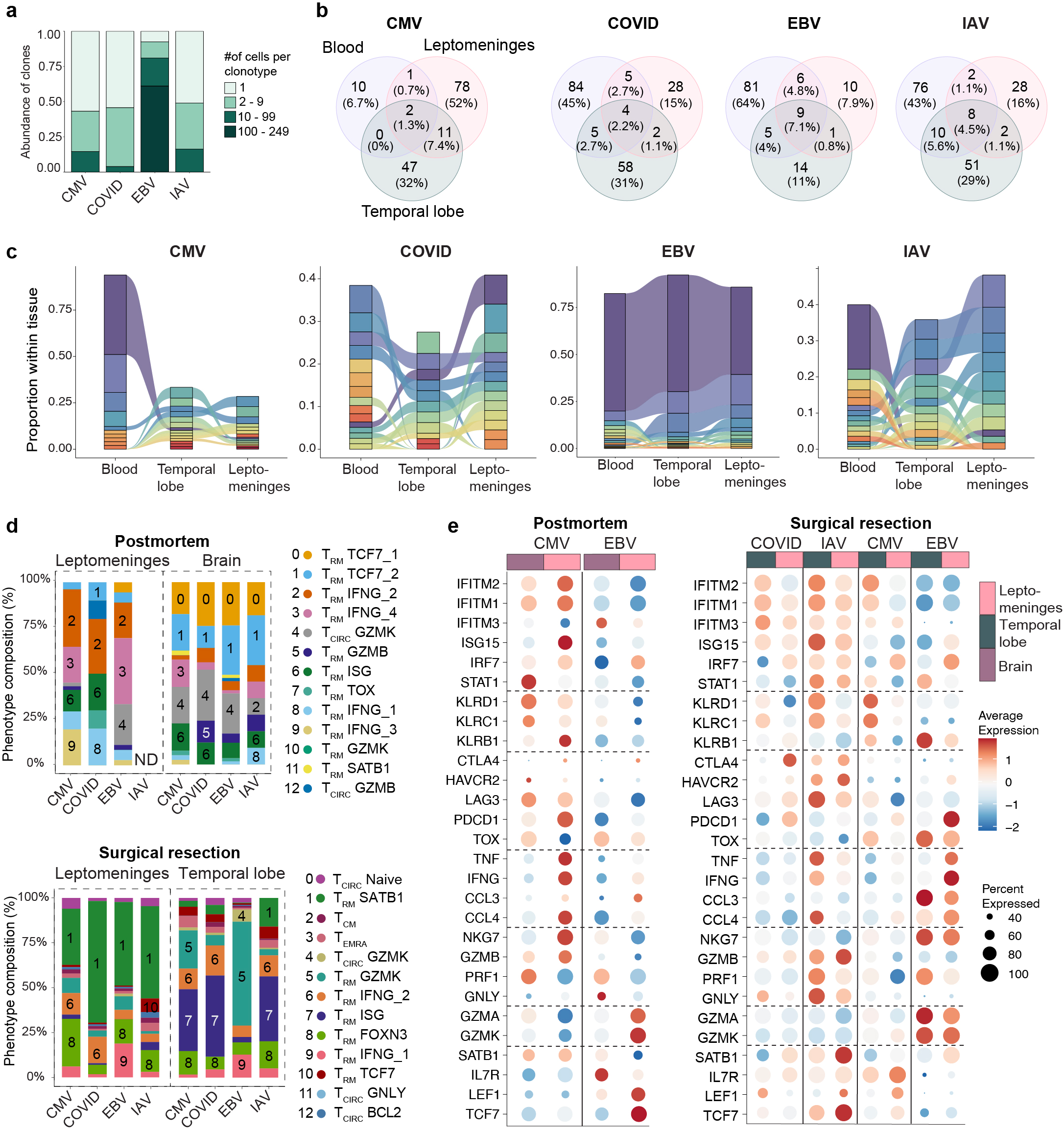
Viral specificity shapes the clonal architecture and phenotype of antigen-specific CNS T_RM_. (**a**) Clone size distribution of tetramer-identified virus-specific CD8^+^ T cells by viral specificity in the SR cohort. (**b**) Venn diagrams showing clonotype overlap between blood, temporal lobe, and leptomeninges for each viral specificity in the SR cohort. Numbers indicate clone counts; percentages indicate fraction of total clones per specificity. (**c**) Alluvial plots showing the phenotypic composition of the top 10 expanded clonotypes across all tissues stratified by viral specificity in the SR cohort. Colors represent clonotype identity; ribbons track individual clonotypes across tissues and are pinched to zero width where a clonotype was not detected. (**d**) Cluster composition of tetramer-identified virus-specific CD8^+^ T cells by viral specificity and tissue compartment in the SR cohort (top)and PM cohort (bottom). Colors represent cluster identity, top clusters per tissues annotated with cluster number. (**e**) Dot plot showing scaled expression of selected genes in tetramer-identified virus-specific CD8^+^ T cells by viral specificity and tissue compartment in the PM (left) and SR right) cohorts. Dot size represents percent of cells expressing each gene; color represents mean scaled expression.

We next examined whether virus-specific CNS T_RM_ differed transcriptionally by antigen specificity in both the SR and PM cohorts. Cluster composition analysis showed that tissue compartment was the primary determinant of virus-specific CD8□T cell phenotypic distribution, with cells from the brain and leptomeningeal tissues occupying largely distinct clusters regardless of antigen specificity, consistent with tissue-associated transcriptional states (**Fig. 5d**). In the PM cohort, brain tissue virus-specific T cells across all four specificities converged predominantly on the TCF7-expressing T_RM_ state (C0, C1), while leptomeningeal cells were distributed across T_RM_-IFNG clusters (C2,3,8,9) **(Fig. 5d)**.

T cells recognizing different antigens generally belonged to the same clusters within each tissue, but when analyzed independently of cluster, they were difference in the expression of specific genes based on viral specificity (**Fig. 5e**). EBV-specific cells were uniquely enriched for *GZMK* and *GZMA* across both tissues and cohorts. In the SR cohort, EBV-specific cells in the leptomeninges additionally showed the highest *IFNG*, *CCL4,* and *TNF* expression of all four specificities. CMV-specific cells in brain compartments were enriched for KLRC1 and KLRD1 across both cohorts, and IAV-specific cells showed a similar KLR signature in brain. COVID and IAV-specific cells were distinguished by elevated *IFITM1, IFITM2*, and *IFITM3* expression in the temporal lobe, consistent with an interferon-stimulated signature. Together, these data show that tissue compartment is the main factor shaping virus-specific CNS CD8□T cell identity, while antigen history adds a secondary, specificity-dependent layer of heterogeneity in gene expression within each tissue compartment.

## DISCUSSION

Here, we find that the human CNS harbors clonally expanded and regionally connected populations of CD8^+^ T_RM_ shaped by common peripheral viral exposures. Combining flow cytometry, scRNAseq and paired TCRseq across two independent human cohorts, we find that anatomic location is the primary determinant of CNS T_RM_ identity and that clonally related cells are distributed across CNS regions yet transcriptionally reflect their local environment. This T_RM_ pool includes T cell specific for EBV, CMV, IAV and COVID, reframing the CNS T cell landscape as, at least in part, a reflection of peripheral immune experiences.

Several foundational studies have shown that the majority of T cells in the mouse brain are resident using functional approaches that assess blood vs tissue localization and migration via intravascular staining and parabiosis^28,32,57^. Defining tissue residency in humans, however, is a challenge, as these functional experiments are not possible and canonical residency markers like CD69 can be upregulated by inflammation^58,59^. This is particularly apparent in the meninges where some reports describe leptomeningeal T cells as resident^7,9^, while others suggest the dura and leptomeninges may be sites of active surveillance by recirculating T cells that egress or inhabit a transition state towards residency^8,60^. Here, we used a residency signature anchored, when available, by matched blood as a circulating reference to assess residency status at the transcriptional level. By this analysis, we found that the majority of T cell clusters across all sampled CNS compartments, including the leptomeninges, scored as resident, with enrichment of *CD69, ITGAE* and *ZNF683* and low/absent *S1PR1* and *SELL* expression. The low TCR repertoire overlap between blood and CNS tissues (Morisita 0.14-0.22), compared with high overlap between temporal lobe and leptomeninges (0.80) provides further evidence for residency rather than continuous surveillance by recirculating T cells.

While T cells across all CNS compartments were largely resident, they were also phenotypically distinct. Leptomeningeal T_RM_ were enriched for immediate-early and NF-κB transcripts (*NR4A1-3,* AP-1 family, *NFKB1*), costimulatory molecules (*ICOS, TNFSF9*), and *IFNG*. Because we and others have not detected corresponding cytokine protein^6^, enrichment of *IFNG* transcript likely reflects a “poised” state where pre-formed cytokine mRNA is held under translational repression until TCR engagement, allowing memory T cells to respond rapidly, as previously described^61^. In contrast, T_RM_ in the brain were comparatively quiescent and enriched for *TGFBR2* (TGFβ receptor) and the inhibitory receptor *KLRC1* (NKG2A). Together, these phenotypes may reflect a more frontline role for meningeal T_RM_ in responding rapidly to infection at the CNS boarder, and a more restrained role for brain T_RM_ where excessive inflammation is poorly tolerated.

This regional imprinting extended to cells of shared clonal origin. We found that expanded clonotypes were broadly shared across CNS regions within individuals, extending prior evidence of brain-leptomeningeal repertoire overlap to multiple anatomically distinct compartments^7^. Interestingly, T cells from the same clonotype consistently adopted the transcriptional state characteristic of the tissue in which it resided, as measured by a regional imprinting score in which each clone served as its own control. Because the distinct tissue signatures were derived exclusively from region-restricted clonotypes, this effect was independent of the tested clones and of sampling depth. This suggests that clonally related cells are not necessarily fated to a fixed CNS T_RM_ program but are likely shaped by local tissue cues. This is consistent with studies showing that different anatomic locations, for example the gut, imprint distinct resident T cell phenotypes^62–64^.

Beyond these compartment-specific transcriptional programs, antigen specificity added a second layer of heterogeneity. Across CNS tissues and cohorts, EBV-specific cells were enriched for *GZMK* and *GZMA*. Interestingly, EBV-specific T cells in the CSF of patients with Alzheimer’s disease also express granzymes A and K^41^. In mice, GZMK-expressing CNS CD8^+^ T cells have been associated with slower tauopathy progression, yet have also been implicated in Alzheimer’s pathology and age-associated cognitive decline^65–67^. While antigen specificity remains unclear in these models, these findings highlight the need to define how granzyme A-and K-expressing CNS T_RM_ influence neurological disease and whether their effects depend on the antigens they recognize.

Other viral specificities also showed distinct transcriptional features. CMV-specific cells were enriched for *KLRC1* and *KLRD1* expression in both cohorts, consistent with the previously described acquisition of CD94 (*KLRD1*) and NKG2A (*KLRC1*) by CMV-specific CD8□ T cells in the blood^68^. In contrast, IAV and COVID-specific cells were generally distinguished by ISG expression, including elevated IFITM2 and IFITM3. IFITM3 has been shown to promote the survival and persistence of lung T_RM_ during influenza infection in mice^69^, raising the possibility that IFITM expression reflects a T cell program imprinted by prior acute infection and/or vaccination which serves to promote long-term T_RM_ survival. Together, these data support a model where the local CNS environment imprints tissue-specific phenotypes that are further influenced by antigen specificity, consistent with patterns observed among antiviral CD8^+^ T cells across non-CNS human tissues^70^.

While we were able to directly characterize rare virus-specific T cell populations in the human CNS, we recognize there are limitations of our study. For one, the small cohort size and heterogeneity in donor backgrounds can limit the generalizability of these findings. Both cohorts were restricted to HLA-A*02-positive donors, restricting assessment of antigen-specific populations across other HLA contexts. The postmortem cohort was predominantly composed of donors over 90 years of age, and age-associated changes in T cell repertoire and phenotype may affect the applicability of these findings to the general population. Postmortem sampling may also introduce transcriptional changes relative to the *in vivo* state, although the reproducibility of major transcriptional programs across postmortem and surgical tissues argues against this being the primary driver of the observed patterns. Finally, the surgical cohort consisted of individuals with drug-refractory epilepsy, which is associated with a pro-inflammatory brain microenvironment in humans^71^ and altered phenotype of brain localized T cells in mouse models^27^, potentially contributing to the virus-specific T cell states observed here.

Despite these limitations, our findings reveal a heterogeneous population of resident CD8□T cells in the human CNS shaped by local tissue and viral specificity. Whether virus-specific T_RM_ in humans provide protective immune surveillance, contribute to pathology, or both, remains an open question. There is abundant evidence in mice that CNS T_RM_ protect against infection^16,18,21,72^, and our findings of clonal expansion and effector-associated gene expression suggest these cells could provide similar protection in humans. Notably, this protection in mice extends to T_RM_ seeded through peripheral vaccination or infection, which carries implications for vaccine design to guard against neurologic infections. These same protective effector functions could become harmful, however, following inappropriate activation or recognition of cross-reactive CNS antigens. This is directly relevant to neurological diseases including multiple sclerosis and Alzheimer’s disease, where EBV-specific clonally expanded CNS CD8□T cells have been found^11,41^. In identifying virus-specific CD8^+^ T cells in the CNS of multiple donors spanning non-diseased brain to Alzheimer’s disease and epilepsy, our work provides a foundation to understand when and how this T_RM_ surveillance is protective and when it becomes a driver of neurological disease.

## METHODS

### Study Cohort

Post-mortem CNS and peripheral tissue specimens were procured from the Department of Pathology and Laboratory Medicine at Dartmouth Hitchcock Medical Center (Lebanon, NH) under an approved institutional protocol. Tissue was collected within a 24-72 hour postmortem interval. Fresh tissue for the surgical resection cohort was obtained from patients undergoing temporal lobe resection for medically refractory epilepsy, enrolled after informed consent under an IRB-approved protocol at Dartmouth Hitchcock Medical Center. Demographic and clinical information for all donors is provided in Table 1.

### Tissue Processing

Human brain tissue was minced into small fragments and enzymatically dissociated in Collagenase Type IV solution supplemented with DNase I at 37°C for 30 minutes for brain samples and 1hr for leptomeninges. Leptomeninges were mechanically dissected from the surface of the brain tissue before processing. Because complete removal from cortical folds and other irregular surfaces was not always possible, residual leptomeningeal tissue may have remained associated with some brain samples. Samples were further mechanically disrupted using a gentleMACS Tissue Dissociator (Miltenyi Biotec) and passed through a 70 μm cell strainer to obtain a single-cell suspension. Myelin debris and fat were removed by density gradient centrifugation using a 44/67% Percoll gradient (Cytiva), and the enriched lymphocyte fraction was washed and resuspended for downstream applications. For samples where immediate processing was not feasible (PM001, 002 003, 007 and 009), tissues were stored at 4°C in MACS Tissue Storage Solution (Miltenyi Biotec) upon receipt for up to ∼12hrs before processing.

### Flow Cytometry

Postmortem tissue-derived cells were stained with a Live/Dead Fixable Blue Dead Cell Stain Kit for UV excitation (Thermo Fisher Scientific, 1:500) for 30 minutes at 4°C to stain non-viable cells. Cells were washed and incubated with a surface antibody panel including CCR7-BV605 (clone G043H7, BioLegend, 1:30), CD69-BUV496 (clone FN50, BD Biosciences, 1:30), CD45RA-BV650 (clone HI100, BioLegend, 1:400), CD103-BV711 (clone Ber-ACT8, BioLegend, 1:100), CD3-Alexa Fluor 700 (clone UCHT1, BioLegend, 1:100), and CD8-APC-Fire 750 (clone SK1, BioLegend, 1:400) for 30 minutes at 4°C. Cells were washed twice with FACS buffer (1× DPBS containing 1% heat-inactivated newborn calf serum) and fixed in 2% paraformaldehyde for 30 minutes prior to acquisition on a Cytek Aurora spectral cytometer. Spectral unmixing and data analysis were performed using SpectroFlo (Cytek) and FlowJo software,r respectively.

### Cell sorting and processing for sequencing

Lymphocytes isolated for sequencing were stained with in-house peptide-MHC tetramers^73^ for 30 minutes at 4°C in PBS supplemented with 2% FBS following Percoll gradient enrichment. All tetramers were combinatorially labelled with APC and barcoded PE (TotalSeq-C0951-54 PE Streptavidin, BioLegend), enabling transcriptional discrimination of each specificity. Peptides used for tetramers were GLCTLVAML (EBV), YLQPRTFLL (SARS-CoV-2), GILGFVFTL (influenza A), and NLVPMVATV (CMV). Cells from all tissues and donors were stained with the four tetramers apart from cells from donor PM001, which was stained with EBV tetramer only for technical reasons. Following tetramer staining, cells were centrifuged and stained with BV711 anti-human CD4 (clone RPA-T4, BioLegend), PE-Cy7 anti-human CD8 (clone SK1, BioLegend), and DAPI (1.5 μg/mL, BioLegend) for 30 minutes at 4°C in PBS with 2% FBS. After two washes, each tissue sample was labelled with a distinct TotalSeq™-C hashtag antibody so that tissues from the same donor could be pooled into a single Chromium lane and demultiplexed computationally. Viable (DAPI-negative) CD8^+^CD4^-^cells were sorted to 80% purity on a Sony SH800S cell sorter. Tetramer-positive cells were sorted separately based on dual labeling of APC and PE-barcoded tetramers. Sorted cells from different tissues were pooled, and combined with the bulk CD8^+^ fraction prior to submission for single-cell RNA sequencing.

### Single-cell RNA-seq and TCR seq analysis

Single-cell gene expression, V(D)J and Feature Barcode libraries were prepared from sorted cells using the 10x Genomics Chromium Next GEM Single Cell 5′ Reagent Kits according to the manufacturer’s protocol. Libraries were pooled and sequenced on an Illumina NextSeq 2000 instrument to a target depth of [30,000] read pairs per cell for gene expression, 5,000 for V(D)J, and 2,000 for Feature Barcode libraries. Gene expression, V(D)J and Feature Barcode libraries were processed together with Cell Ranger multi (10x Genomics, v9.0.0) against the GRCh38 reference (refdata-gex-GRCh38-2024-A) and the human V(D)J reference (refdata-cellranger-vdj-GRCh38-alts-ensembl-7.1.0), using a feature reference CSV specifying the tetramer and hashtag barcode sequences.

Downstream analysis was performed in R (v4.3) using Seurat (v5.0) and tidyverse packages. Samples were demultiplexed using HTODemux. Cells with normalized *CD8A* and *CD8B* expression >0.01 were retained, and cells with <500 or >4,000 detected genes or >10% mitochondrial transcripts were excluded. Data were normalized using SCTransform with mitochondrial transcript proportion regressed out. TCR-encoding genes were excluded from highly variable features before principal component analysis. Samples were integrated using reciprocal PCA, and UMAP embeddings were generated using the first 30 principal components. Gene expression was imputed using adaptively thresholded low-rank approximation (ALRA)^74^ for downstream analysis. TCR repertoire analyses were performed using scRepertoire. Clonotypes were defined by CDR3 nucleotide sequence and V(D)J gene usage of recovered TCRα and TCRβ chains and were indexed by patient to maintain donor-specific clonotype assignments.

### Barcoded tetramer identification

Antigen-specific cells were first identified based on barcode expression using thresholds at points of bimodal separation between positive and negative populations. Cells exceeding the threshold for more than one specificity were flagged as potential doublets and excluded from downstream analyses. Remaining cells were assigned a preliminary tetramer identity based on each specificity. Tetramer calls were further refined using paired TCR clonotype information. Tetramer-positive barcodes were intersected with cells containing paired TCR sequencing data, and associated clonotypes (CTstrict) were identified using scRepertoire. All cells across the dataset sharing these clonotypes were then retrieved, including cells that did not individually exceed the tetramer threshold, based on the assumption that cells sharing an identical TCR clonotype have the same antigen specificity. To further reduce false-positive assignments, a clone-level filter was applied where clonotypes were retained only when the number of cells with the cognate tetramer signal exceeded the number of unlabeled cells within that clonotype.

### MixTCRpred

For each cell, paired TCRα and TCRβ sequences were extracted from VDJ annotations, and the highest-scoring αβ pair was retained when multiple chains were present. TCR sequences were scored against four HLA-A*02:01–restricted epitope models (GILGFVFTL, YLQPRTFLL, GLCTLVAML, NLVPMVATV). As a result, each TCR sequence will have four scores, where a score is the %rank of the query relative to the binding prediction of background TCRs. These background TCRs were integrated in the MixTCRpred package. The true binders were expected to have the %rank less than 0.1^51^. Therefore, we retained the TCR sequences with exactly one %rank <0.1 among the four scores, and the best scored epitope is the predicted epitope. Predicted epitope assignments were mapped back to single-cell transcriptomic data using cell barcodes and added to the Seurat metadata for downstream analysis. Predicted epitope assignments were compared with tetramer-based specificity annotations to assess concordance between approaches.

### VDJdb matching

TCR sequences were compared with the curated VDJdb database of antigen-specific TCRs. Matches were assigned based exact CDR3 amino acid sequence identity and available chain information. Paired TRA–TRB matches were prioritized when available, whereas exact TRB CDR3 matches were used for descriptive antigen-specificity assignments when paired-chain matches were not recovered. Viral specificity was assigned according to the antigen species annotated in VDJdb. For visualization, figures show exact TRB CDR3 matches corresponding to the four viral specificities assessed by tetramer staining (EBV, CMV, IAV and COVID). All VDJdb TRB matches identified in the dataset are reported in Supplementary Table 2, and paired exact TRA–TRB matches are reported separately in Supplementary Table 3.

### Tissue residency scoring

T_RM_ upregulated and -downregulated gene sets were taken from Burn et al.^53^ and filtered to genes detected in our dataset. Module scores were computed per cell with UCell on the SCTransform-normalized assay. A residency index was calculated as the difference between resident and circulating scores (resident minus circulating), with higher values indicating a more resident-like transcriptional state. In the SR cohort with matched blood and tissue, blood-derived cells were used as a circulating reference to anchor classification thresholds. For PM dataset, cells were classified based on relative dominance of the two programs: resident-like when resident scores exceeded circulating scores and circulating when circulating scores exceeded resident scores. Cluster annotations were assigned based on the dominant per-cell state within each cluster.

### Differential expression and cross-dataset overlap

Differentially expressed genes between brain and leptomeningeal T cells were identified separately in each dataset using FindMarkers(Seurat) with a Wilcoxon rank-sum test, retaining genes with Bonferroni-adjusted *P* < 0.05 and classifying them by the sign of the average log2 fold change. Genes significant in both datasets in the same direction were taken as a reproducible signature and ranked by mean adjusted *P* across datasets, with mean absolute log2 fold change as a tiebreaker.

### Cross-cohort cluster similarity analysis

To compare transcriptional programs across independently processed cohorts, we generated pseudobulk expression profiles for each T cell cluster within each cohort by aggregating raw counts across all cells belonging to a cluster. Counts were normalized using Trimmed Mean of M-values (TMM) normalization to account for differences in library size and compositional bias and transformed to log counts per million (logCPM). Replicate-level pseudobulks were averaged to obtain a single cluster centroid expression profile for each cluster. Similarity between clusters across cohorts was quantified using Spearman correlation computed across genes shared between both datasets, generating a cluster-by-cluster similarity matrix.

### Digestion-associated gene PCA

To test whether tissue-associated transcriptional differences were driven by enzymatic digestion artifacts, pseudobulk profiles were generated by summing raw RNA counts per donor-tissue combination. Collagenase-associated genes were defined from the kidney collagenase-versus-cold-active protease comparison reported by Crowl et al.^56^, retaining genes with mean expression above threshold and greater than 1.5-fold higher expression following collagenase digestion, with mouse genes mapped to human orthologs. The 2,000 most variable genes were selected after TMM normalization and log2 CPM transformation, and all detected collagenase-associated genes were added to this feature set. PCA was performed on centered and scaled expression values before and after removal of collagenase-associated genes.

### Repertoire overlap between tissues

Clonal overlap between tissues was quantified within each patient separately. Cells lacking a clonotype call or tissue annotation were excluded. For each patient, pairwise overlap between every pair of tissues was computed using two complementary indices: the Morisita–Horn index, calculated on clonotype cell counts as C = 2·Σ(x_i·y_i) / [(D_x + D_y)·X·Y], where X and Y are the total cells per tissue and D = Σx_i²/X²; and the Jaccard index, calculated on the unordered sets of unique clonotypes as |A∩B| / |A∪B|. Tissue pairs for which a patient had no cells in one or both tissues were set to NA. Per-patient matrices were then averaged element-wise across patients.

### Regional imprinting of shared clonotypes

To test whether cells of a clonotype present in more than one CNS region adopt different states by location, we grouped tissues into two compartments— brain parenchyma and leptomeninges (blood excluded)— and compared cells within each clonotype across compartments. To avoid circularity, tissue-specific signatures were derived only from cells belonging to clonotypes confined to a single compartment. Within these cells, we identified genes differentially expressed between tissue compartments (Wilcoxon test, Benjamini-Hochberg correction), restricting to genes detected in ≥20% of cells and removing non-T-cell, ribosomal, mitochondrial, and stress/dissociation-associated genes. The top genes in each direction formed a brain and a leptomeningeal signature. Each cell was scored for both signatures with UCell, and a polarization score δ was defined per cell as brain minus leptomeningeal score. Shared clonotypes were those with cells in both compartments within one donor. For each, a regional imprinting score was calculated as mean δ of its brain cells minus mean δ of its leptomeningeal cells. The score is positive when a clonotype’s brain cells are more brain-like than its leptomeningeal cells, negative when the reverse holds, and near zero when the two are similar. We also examined the mean δ of each clonotype separately per compartment, since the score measures the difference between compartments rather than the absolute position of each.

### Cell population definitions and nomenclature

Definition of T cell nomenclature used in this paper is outlined here and was determined by either flow cytometry phenotyping or by scRNA-seq cluster identity^75^. **Flow cytometry-defined cells:** CD8^+^ T cells were classified by CCR7 and CD45RA expression as naive (CCR7^+^CD45RA^+^) ^76^. CD69 and CD103 were used as markers of tissue residency, with CD69^+^ and CD69^+^CD103^+^ CNS CD8^+^ T cells defined as resident^59^. **scRNA-seq-defined cells:** CNS CD8^+^ T cell clusters were first classified as circulating (T_CIRC_ here, also known as TDM) or tissue resident (T_RM_ here, also known as TD_R_M) using module scores derived from published circulation and tissue-residency gene signatures, together with canonical markers such as *S1PR1* and *SELL* for circulating cells and *CXCR6*, *ZNF683*, and *ITGAE* for resident cells. T_CIRC_ clusters were then further annotated follows: naive cells were defined by expression of CCR7, SELL, TCF7 and LEF1. The BCL2+ memory-like population was defined by BCL2 together with CD28, FAS and BCL2 and reduced expression of naive-associated genes. Central memory cells (T_CM_ here, also known as TSM) were defined by retention of the naive-associated program together with CD27, CXCR3 and LTB. Effector memory cells (T_EM_ or T_EMRA_ here, also known as TD_W_M) were defined by NKG7, PRF1, GNLY and GZMB, while T_EMRA_ cells were defined by expression of CX3CR1, B3GAT1, FCGR3A and FGFBP2. A subset of T_EM_ that were high for GZMK were labeled as such. T_RM_ clusters were named by their main cluster-defining genes or functional programs, including stem/progenitor-like, GZMK, SATB1, FOXN3, IFNG/cytokine and ISG states.

## Acknowledgments

We thank the patients and donor families whose contributions made this study possible. We thank Lauren J. Sinks and Beverly A. Hills for assistance in patient recruitment and coordination. We thank the Dartmouth Cancer Center Shared Resources and bioMT core at Dartmouth College for their contributions to this work. We thank Gary Ward at DartLab, Immune Monitoring and Flow Cytometry Shared Resource for help with FACS and flow cytometery.

## Author contributions

Conceptualization: HND, PCR

Methodology: HND, TGS, LS, PCR

Analysis: HND, TGS, LS

Investigation: HND, SCM, TGS, SAK, TC, MAF, JFI, SY

Visualization: HND, PCR

Funding acquisition: PCR

Project administration: PCR

Resources: FWK, GJZ, CPL, JH, AGJS, MJT, PCR

Supervision: PCR

Writing – original draft: HND, PCR

Writing – review & editing: HND, PCR

## Data and materials availability

All sequencing data generated in this study will be deposited in GEO. Custom analysis code will be made publicly available upon publication

