## Supplemental Figures for "Common viral infections seed regionally distinct resident memory T cells in the human CNS"

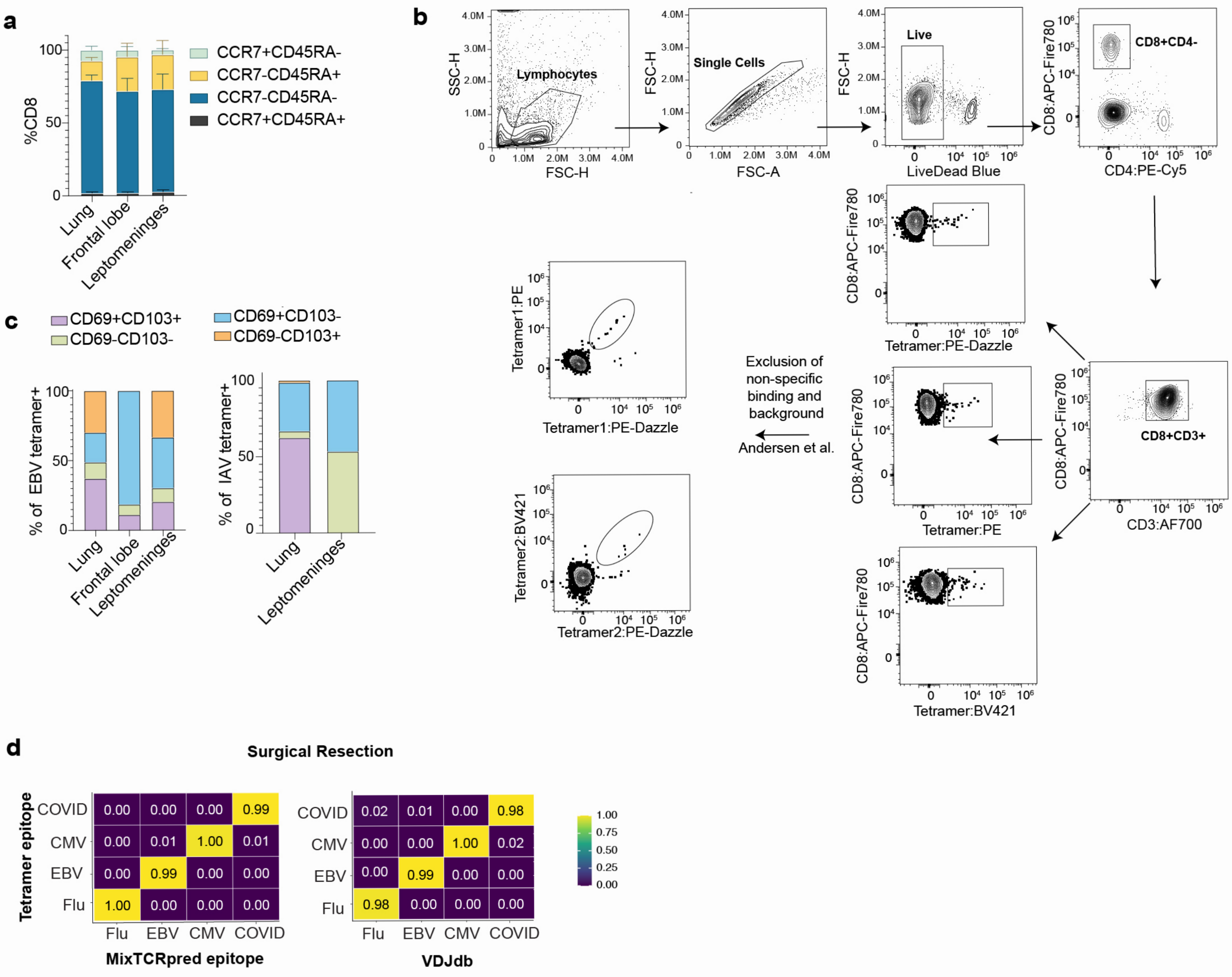


**Supp Data Figure 1. Identification of virus-specific CD8+ T cells.**
(**a**) CCR7 and CD45RA co-expression among $CD8^+$ T cells across tissues (n=5; bars represent mean ± SEM). (**b**) Gating strategy for combinatorial tetramer staining. (**c**) CD69 and CD103 co-expression among EBV and IAV tetramer positive cells. Data points representing fewer than 10 tetramer positive cells were excluded (bars represent mean ± SEM). (**d**) Concordance between MixTCR- (left) and VDJdb (right) predicted tetramer defined viral specificity. Fractions are normalized within each MixTCR or VDJdb-predicted specificity.

**a**

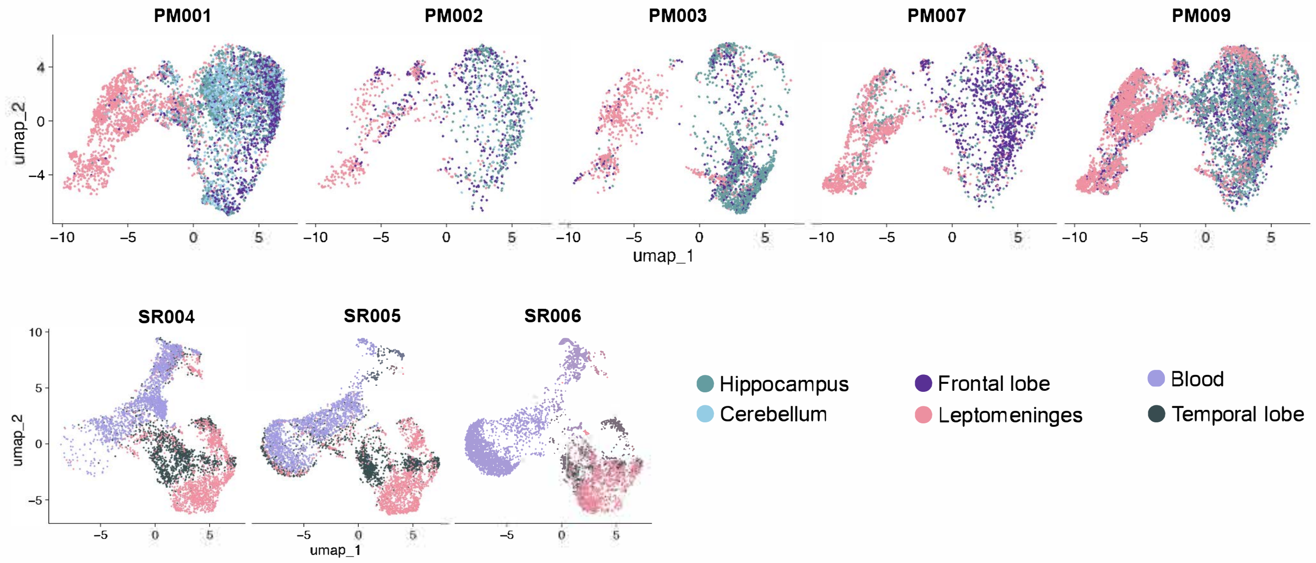


**Supp Data Figure 2. Donor level transcriptional variation of CNS CD8⁺ T cells.** (**a**) UMAP of CD8⁺ T cells colored by tissue of origin, faceted by individual donor for PM cohort (top) and SR cohort (bottom).

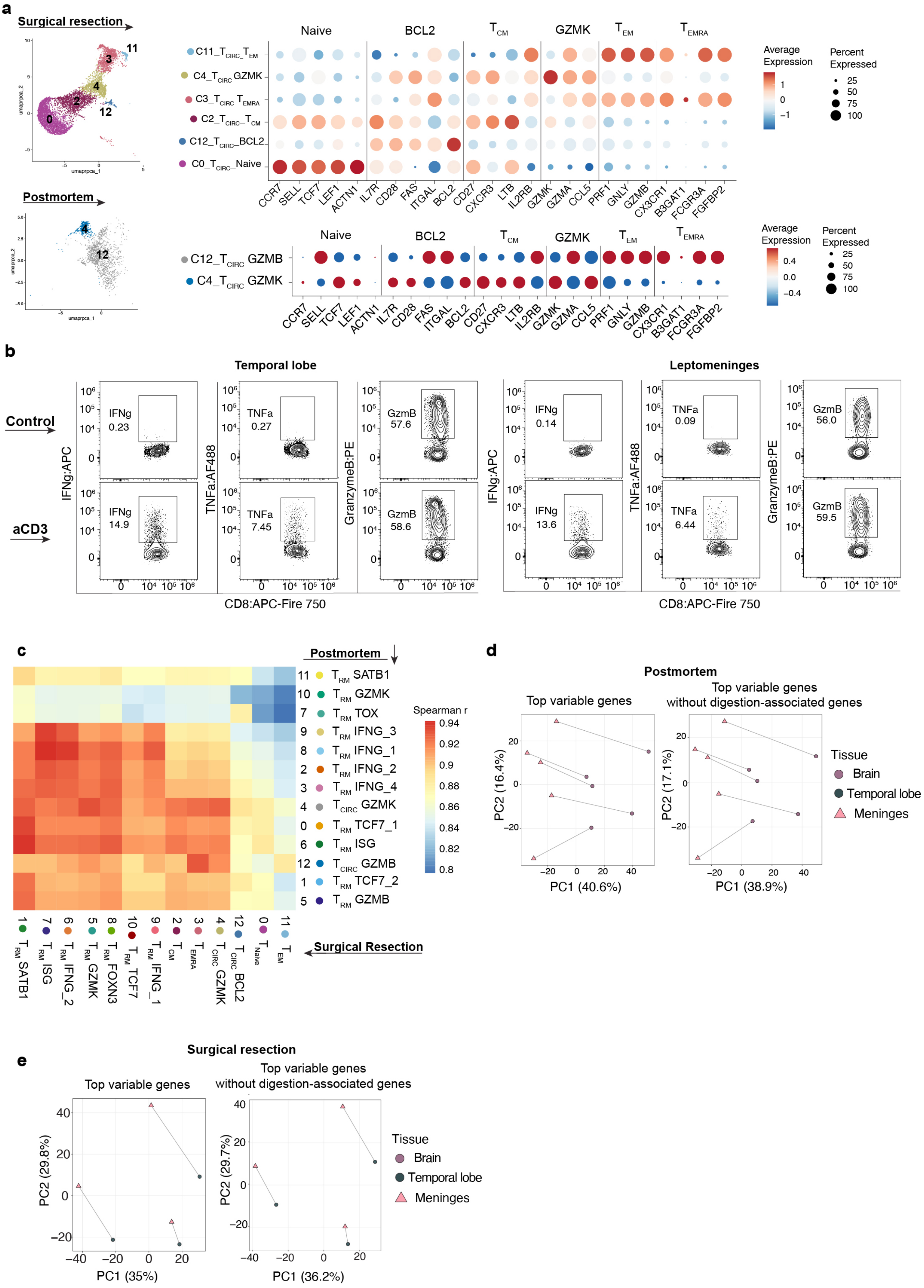
a
Surgical resection
Naive
BCL2
TCM
GZMK
TEM
TEMRA
C11_TCIRC_TEM
C4_TCIRC GZMK
C3_TCIRC TEMRA
C2_TCIRC–TCM
C12_TCIRC–BCL2
C0_TCIRC–Naive
CCR7 SELL TCF7 LEF1 ACTN1 IL7R CD28 FAS ITGAL BCL2 CD27 CXCR3 LTB IL2RB GZMK GZMA CCL5 PRF1 GNLY GZMB CX3CR1 B3GAT1 FCGR3A FGFBP2
Average Expression
Percent Expressed
25
50
75
100
Postmortem
C12_TCIRC GZMB
C4_TCIRC GZMK
umaprpca_1
umaprpca_2
b
Temporal lobe
Leptomeninges
Control
aCD3
IFNg:APC
TNFa:AF488
GranzymeB:PE
IFNg 0.23
TNFa 0.27
GzmB 57.6
IFNg 14.9
TNFa 7.45
GzmB 58.6
IFNg 0.14
TNFa 0.09
GzmB 56.0
IFNg 13.6
TNFa 6.44
GzmB 59.5
CD8:APC-Fire 750
c
Postmortem
11 TRM SATB1
10 TRM GZMK
7 TRM TOX
9 TRM IFNG_3
8 TRM IFNG_1
2 TRM IFNG_2
3 TRM IFNG_4
4 TCIRC GZMK
0 TRM TCF7_1
6 TRM ISG
12 TCIRC GZMB
1 TRM TCF7_2
5 TRM GZMB
Spearman r
1 TRM SATB1
7 TRM ISG
6 TRM IFNG_2
5 TRM GZMK
8 TRM FOXN3
10 TRM TCF7
9 TRM IFNG_1
2 TCM
3 TEMRA
4 TCIRC GZMK
12 TCIRC BCL2
0 TNaive
11 TEM
Surgical Resection
d
Postmortem
Top variable genes
Top variable genes without digestion-associated genes
PC1 (40.6%)
PC2 (16.4%)
PC1 (38.9%)
PC2 (17.1%)
Tissue
Brain
Temporal lobe
Meninges
e
Surgical resection
Top variable genes
Top variable genes without digestion-associated genes
PC1 (35%)
PC2 (29.8%)
PC1 (36.2%)
PC2 (29.7%)
Tissue
Brain
Temporal lobe
Meninges

**Supp Data Figure 3. Circulating CD8$^+$ T cell clusters and transcriptional quality controls.** (**a**) UMAP of circulating CD8$^+$ T cell clusters from SR cohort (top) and PM cohort (bottom), with dot plots showing scaled expression of selected genes used to define cluster identities. Dot size represents percent of cells expressing each gene; color represents mean scaled expression. (**b**) Intracellular cytokine staining for granzyme B, IFNγ and TNFα in an unstimulated (top row) and anti-CD3 stimulated (bottom row) CD8+ T cells from temporal lobe and leptomeninges. Gated on live CD8+ T cells; n=1. (**c**) PCA analysis of pseudobulk transcriptomes before (left) and after (right) removal of collagenase digestion-associated genes. Each point represents one donor-tissue pseudobulk profile; lines connect matched tissues from the same donor. (**d**) Pseudobulk Spearman correlation heatmap showing pairwise concordance of gene expression between cluster from PM and SR cohorts.

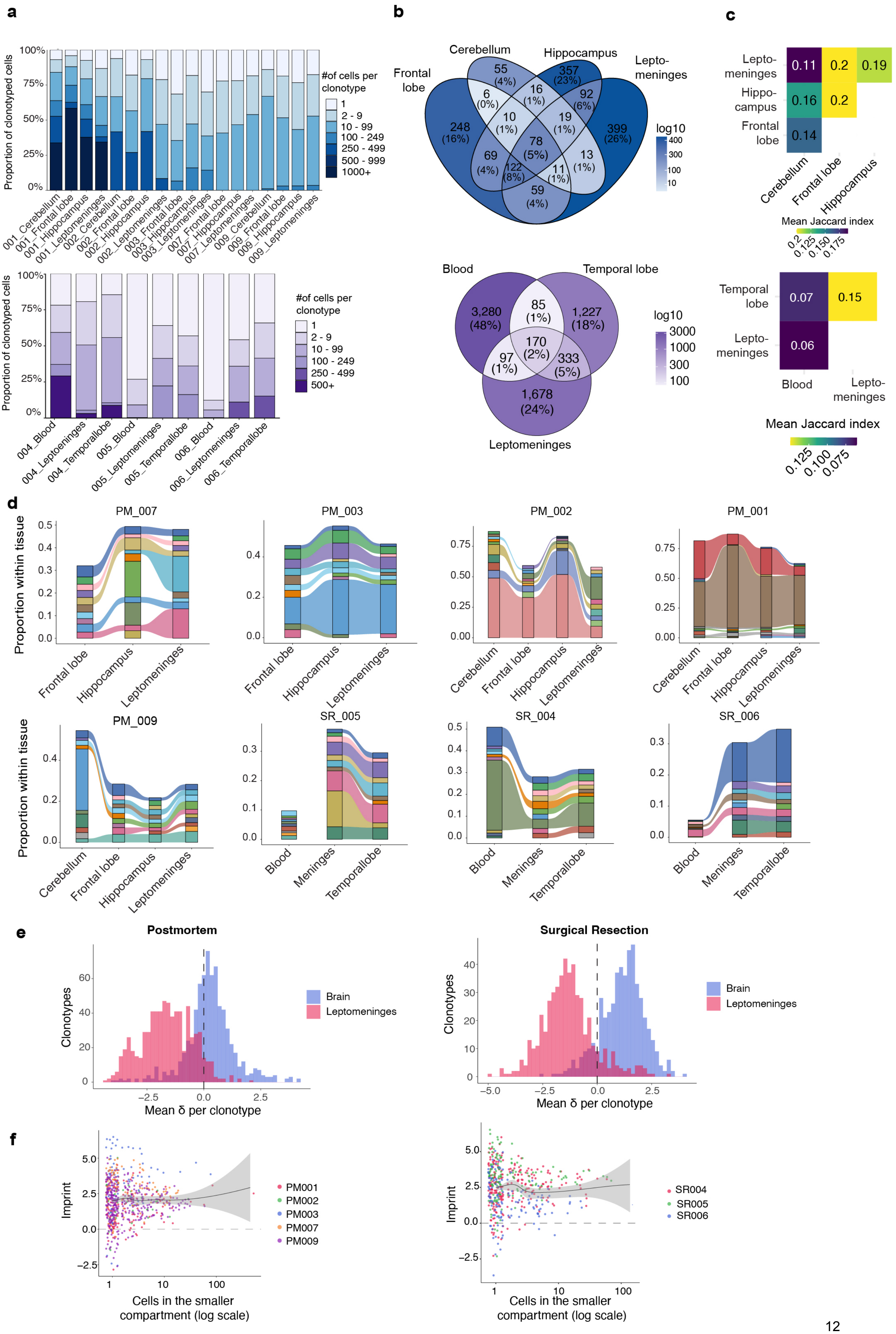
a
100%
75%
50%
25%
0%
Proportion of clonotyped cells
#of cells per clonotype
1
2 - 9
10 - 99
100 - 249
250 - 499
500 - 999
1000+
001_Cerebellum
001_Frontal lobe
001_Hippocampus
001_Leptomeninges
002_Cerebellum
002_Frontal lobe
002_Hippocampus
002_Leptomeninges
003_Frontal lobe
003_Hippocampus
003_Leptomeninges
007_Frontal lobe
007_Hippocampus
007_Leptomeninges
009_Cerebellum
009_Frontal lobe
009_Hippocampus
009_Leptomeninges
500+
004_Blood
004_Leptoeninges
004_Temporallobe
005_Blood
005_Leptomeninges
005_Temporallobe
006_Blood
006_Leptomeninges
006_Temporallobe
b
Cerebellum
Hippocampus
Frontal lobe
Lepto-meninges
55 (4%)
357 (23%)
6 (0%)
16 (1%)
92 (6%)
248 (16%)
10 (1%)
19 (1%)
399 (26%)
78 (5%)
69 (4%)
13 (1%)
122 (8%)
11 (1%)
59 (4%)
log10
400
300
100
10
Blood
Temporal lobe
3,280 (48%)
85 (1%)
1,227 (18%)
170 (2%)
97 (1%)
333 (5%)
1,678 (24%)
Leptomeninges
log10
3000
1000
300
100
c
Lepto-meninges
Hippo-campus
Frontal lobe
0.11
0.2
0.19
0.16
0.2
0.14
Cerebellum
Frontal lobe
Hippocampus
Mean Jaccard index
0.2
0.125
0.150
0.175
Temporal lobe
Lepto-meninges
0.07
0.15
0.06
Blood
Lepto-meninges
Mean Jaccard index
0.125
0.100
0.075
d
Proportion within tissue
PM_007
PM_003
PM_002
PM_001
Frontal lobe
Hippocampus
Leptomeninges
Cerebellum
PM_009
SR_005
SR_004
SR_006
Blood
Meninges
Temporallobe
e
Postmortem
Surgical Resection
Clonotypes
Mean δ per clonotype
Brain
Leptomeninges
f
Imprint
Cells in the smaller compartment (log scale)
PM001
PM002
PM003
PM007
PM009
SR004
SR005
SR006

**Supp Data Figure 4. Repertoire structure and clonal sharing across CNS regions.**(**a**) Proportion of clonotyped $CD8^+$ T cells in patient-wide clone-size bins across tissues, shown by patient for PM (top) and SR (bottom). (**b**) Venn diagram showing overlap of T cell clonotypes across tissues or PM (top) and SR (bottom). (**c**) Jaccard overlap index heatmap showing pairwise clonal similarity between tissues in the PM cohort (top) and SR cohort (bottom). Values represent the mean Jaccard index across donors. (**d**) Alluvial plots showing the top 10 expanded clonotypes per donor, tracked across tissue compartments. Ribbon width is proportional to the number of cells belonging to each clonotype in each tissue. Colors represent individual clonotypes. (**e**) Polarization score (δ) of shared clonotypes by tissue compartment. For each clonotype with cells in both compartments, the mean δ of its parenchymal cells and of its meningeal cells is shown. (**f**) Regional imprinting score per shared clonotype versus the number of cells in its less-represented compartment (log scale); each point is one clonotype, colored by donor. Line, LOESS fit; shaded band, 95% confidence interval; dashed line, no imprinting. PM (left), SR (right).
